# NanoPro – a Framework for Probiotically-Guided Nanovector Therapy

**DOI:** 10.64898/2026.10.28.762486

**Authors:** Alexander Michels, Azra Haseki, Myra Ashraf, Audrey Lam, Kailyn R. Grant, Amy Lu, Rosa L. Vincent, Jongwon Im, Zakary S. Singer, Edward R. Ballister, Tal Danino

## Abstract

The “antigen problem”, i.e. the disconnect between the broad spectrum of therapeutic modes enabled by the range of viral and synthetic nanoparticles for which workable receptor-targeting architectures exist and the relative dearth of addressable target receptors on most tumors, is a long-standing challenge in targeted tumor nanotherapy. Here we provide *in vitro*-proof-of-principle for NanoPro, a platform for guiding nanoscale therapeutics to otherwise antigenically indistinct tumors via probiotically-released tumor cell-binding tags, ProTags. These consist of an orthogonal antigen (GFP or ALFA) fused to the heparin-binding domain of placenta growth factor-2 and serve as attachment receptors for ProTag-targeted nanovectors. ProTag release from *Escherichia coli* Nissle 1917 was facilitated by deletion of the membrane-tethering *lpp* gene, which increased ProTag supernatant concentrations 16.5–fold compared to the parental strain without impacting viability. HBD-dependent decoration with ProTag in bacterial supernatant followed by treatment with receptor-targeted aGFP- or aALFA-lentiviral vectors resulted in majority transduction of MC-38 and MDA-MB-468 tumor cell lines with antigen specificities of 96.5% and 98.5%, respectively, depending on the choice of nanobody. This ProTag-mediated interaction was notably robust, with selective transduction still observed at 625–fold supernatant dilution, warranting further exploration of the utility of the NanoPro system for tumor-restricted delivery of nanotherapeutics.

## INTRODUCTION

Receptor-targeting is a technique of rational design in which binders specific for receptors-of-interest are introduced into the particle structures of viral and synthetic nanoscale vectors to alter their tropism. Driven in large part by the needs of the next generation of *in vivo* gene therapies, the last ten years of innovation in receptor-targeting have demonstrated the broad applicability of its principles, yielding highly selective targeted architectures for all major classes of particles^1,2^. In light of the considerable range of therapeutic modes enabled by such a diversity of particle configurations, the direct antitumor application of targeted nanoscale vectors, beyond their use for *in vivo* CAR T cell generation, should be explored.

A major hurdle for direct applications so far has been the relative dearth of actionable antigens in most tumors. Recent work done in our lab demonstrated that this discrepancy can be circumvented by exploiting the ability of bacteria to colonize tumors in an antigen-independent manner: Probiotic *Escherichia coli* Nissle 1917 were engineered to release ProTag, an adaptor protein consisting of GFP as an orthogonal antigen fused to an extracellular-matrix binding domain, which decorated tumor cells both *in vitro* and *in vivo* and facilitated tumor clearance by engaging aGFP-CAR-T cells^3^.

The orthogonal antigenic interface created by the ProTag might enable NanoPro, a system of therapy using probiotically-guided nanovectors which deliver high-impact modalities such as oncolytic viruses and mRNA-LNPs in a spatially controlled, tumor-restricted manner (**Fig. 1**). Towards such a system, in this work we engineer and characterize privileged interactions between ProTag-decorated mammalian cells and Protag-specific vectors which selectively bridge the micro- and nanoscales.

**Figure 1:**
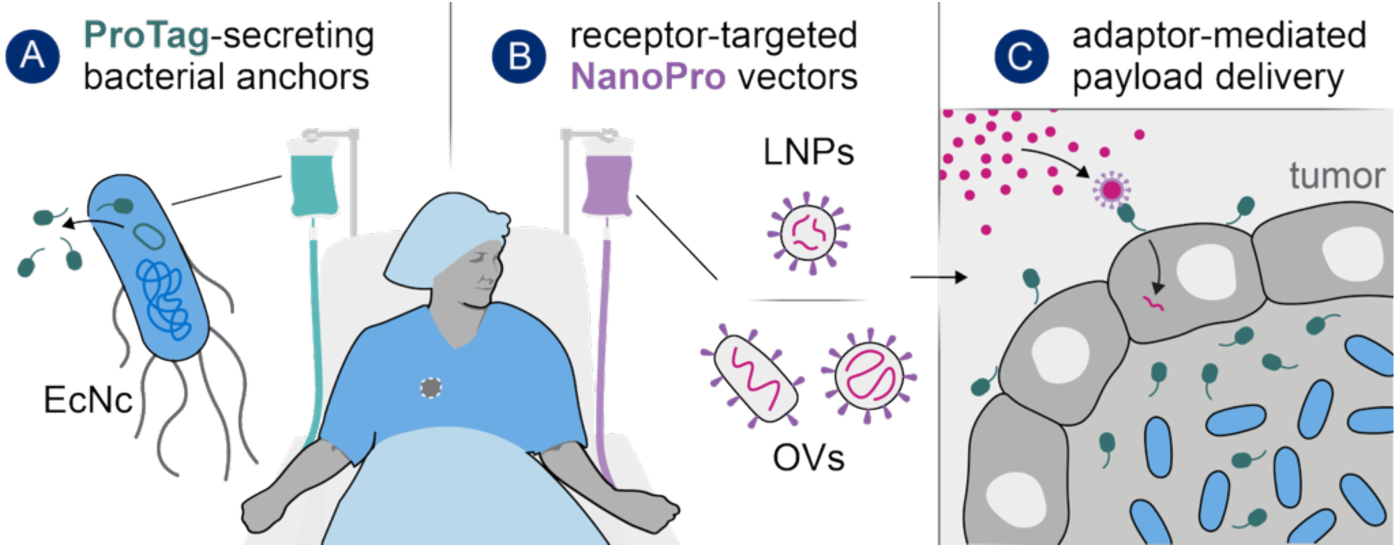
Bacterially-guided tumor drug delivery with targeted nanoscale vectors. Schematic overview of the envisioned system: A NanoPro treatment regime would involve, first (**A**) the administration of an engineered bacterial strain e.g. based on *E. coli* Nissle 1917 (EcNc). These colonize solid tumors and secrete the ProTag adaptor protein (green). (**B**) Subsequently administered NanoPro vectors –such as mRNA-lipid nanoparticles (LNPs) or oncolytic viruses (OVs)– would then (**C**) deliver their payload to ProTag-decorated tumor cells. Towards such a system, we here provide proof-of-principle for bacterially mediated reporter transfer by NanoPro-LVs.

## RESULTS

One modality-of-interest for the NanoPro system are oncolytic viruses, for which probiotically-released, tumor-restricted orthogonal entry receptors may help address the efficacy:safety tradeoff central to their clinical development and allow applications beyond self-experimentation^4^. For platform development, we designed ProTag-specific lentiviral vectors (NanoPro-LVs) transferring an mScarlet-encoding reporter cassette as a surrogate for ProTag-specific oncolytic viruses. NanoPro-LVs feature a targeting pseudotype consisting of i) VSV G protein, mutated to ablate its natural LDLR-tropism and increase serum/thermal stability^5^; as well as ii) a chimeric Display protein^6^ constructed using segments derived from human IgG1 (mutated to disrupt interactions with FcγRs^7,8^), CD28, and HIV-1 gp41 presenting nanobodies cognate for the orthogonal antigens GFP or ALFA-Tag^9^ on the particle surface (**Fig. 2AB**). To enable evaluation of this architecture independently of the bacterial decoration aspect of the proposed interface, we generated genetically decorated derivatives of the MDA-MB-468 human breast cancer cell line –ProMimic– which mimic ProTag decoration by displaying GFP or ALFA on their surface via hCD8a-derived anchors (**Fig. 2C**).

**Figure 2:**
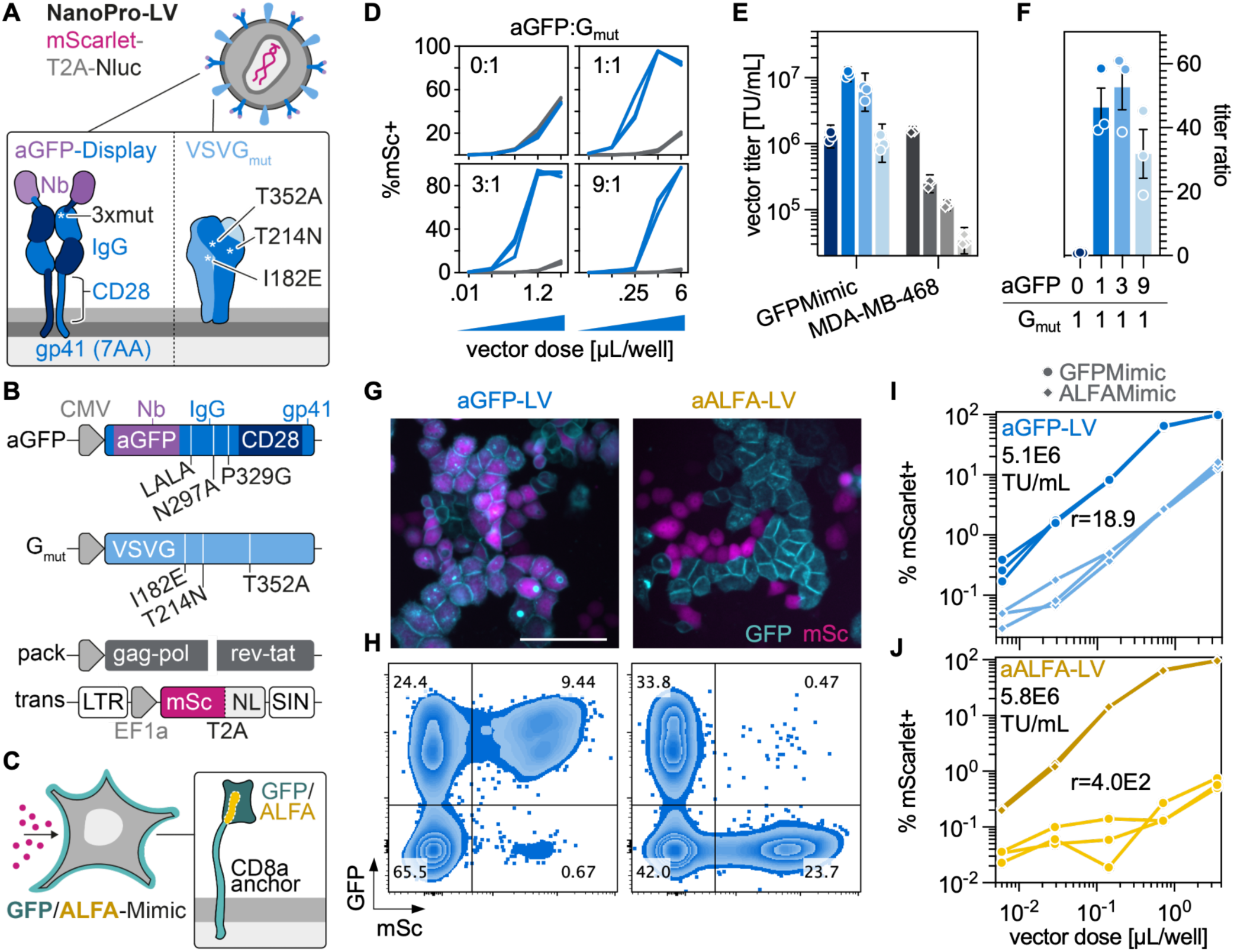
NanoPro-LVs efficiently and selectively transduce tumor cells via orthogonal antigens. (**A-B**) Architecture of NanoPro-lentiviral vectors (NanoPro-LVs). The targeting pseudotype employed here uses a i) chimeric display proteins composed of aGFP or aALFA nanobody plus domains from human IgG1 Fc (mutated to ablate FcR interactions, LALA: L234A/L235A) and CD28 as well as the seven C-terminal amino acids of HIV-1 gp41; and ii) triple-mutated vesicular stomatitis virus G protein (VSVG_mut_)^5–8^. All LV stocks used here were produced with second-generation packaging plasmids as well as transfer plasmids with self-inactivating (SIN) 3’ long terminal repeats (LTRs). MSc: mScarlet; NL: nanoluciferase (**C**) MDA-MB-468 cells displaying GFP^3^ or ALFA^9^ on their surface via hCD8a-derived anchors were used as a surrogate for ProTag-bound tumor cells in transduction assays. (**D-F**) Determining optimal plasmid ratios for production of NanoPro-LVs. GFP-Mimic (blue) or parental MDA-MB-468 cells (gray) were treated with aGFP-LVs produced with the indicated weight ratios of aGFP-Display- to VSVG_mut_-encoding envelope plasmids (**D**) mScarlet-positivities among singlets as determined by flow cytometry four days after vector treatment (4 dpt). Traces represent data from n=3 technical replicates per condition. (**E**) Titers of aGFP-LVs with the previously specified aGFP:G_mut_ ratios on target-(GFP-Mimic) and non-target (MDA-MB-468) cells. Lighter bars indicate higher proportions of aGFP-Display plasmids. (**F**) Ratios of on-over off-target titers. Symbols represent data from n=3 technical replicates, bars & error bars represent geometric means ± 95% CIs for (E) and means ± SEMs for (F). (**G-J**) Assessing the target-specificity of NanoPro-LV transduction. One-to-one cocultures of ALFA- and GFP-Mimic cells were treated with 5 µL/6- well of 3:1 Display:G_mut_ aGFP- or aALFA-LVs and analyzed by (**G**) fluorescence microscopy and (**H**) flow cytometry 3 dpt. Scale bar represents 100 µm. Micrographs (G) and cytometry plots (H) are from two sequential runs with different vector batches. Additionally, aGFP- (**I**) and aALFA-LVs (**J**) were titrated on ALFA- and GFP-Mimic monocultures, and the on-over off-target titer ratios (r) were calculated. Shown are mScarlet-positives among singlet cells v LV doses (per 96-well) 4 dpt. Traces represent data from n=3 technical replicates per condition.

The respective contributions of the aGFP-Display protein and VSVG_mut_ to particle behaviour became apparent when LV stocks produced with different ratios of the plasmids encoding them were titrated on either GFPMimic cells or the parental line (**Fig. 2D**). While VSVG_mut_ alone mediated mScarlet gene transfer at similar efficiency on both GFPMimic and parental cells, transfer rate to non-target cells decreased with the proportion of G_mut_-encoding plasmid in the envelope plasmid mix used during particle production. Conversely, introduction of aGFP-Display protein at aGFP:G_mut_ ratios of 1:1 and 3:1 increased on-target titers by 8.8- and 4.8-fold, respectively, relative to that of aGFP-less particles. A 9:1 ratio did not result in higher on-target titers than aGFP-less LVs, perhaps because of an increased proportion of G_mut_-less particles as fewer production cells receive at least one copy of G_mut_ plasmid (**Fig. 2E**). Crucially, all aGFP-containing configurations were markedly selective for their target antigen, displaying on-over off-target titer ratios ranging from 32 – 53, with the latter observed for an aGFP:G_mut_ ratio of 3:1 (**Fig. 2F**). Based on this, the 3:1 ratio was used for all following LV productions. Orthogonal target specificity was not restricted to GFP-targeting, and mutually exclusive: When cocultures of ALFA- and GFPMimic cells were treated with either aALFA- or aGFP-LV, 98.1% and 93.4% of transfer signal, respectively, originated from the target-expressing compartment (**Fig. 2GH**). A higher target-specificity of aALFA-relative to aGFP-LVs was also observed in titrations on ALFA- and GFP-Mimic monocultures, where aALFA-LVs displayed a 400-fold versus the 19-fold titer ratio obtained with aGFP-LVs (corresponding to 99.75% and 94.7% specificity, respectively; **Fig. 2IJ**), suggesting off-target activity of the aGFP nanobody. Consistent with this, binding affinities of *K_D_ =* 0.59 nM and *K_D_ =* 26 pM have been reported for the used aGFP-^10^ and aALFA-Nbs^9^, respectively, albeit using different approaches.

Having demonstrated the ability of NanoPro-LVs to transduce tumor cells via genetically surface-displayed orthogonal antigens, we next explored whether such selective interaction could also be mediated by probiotically-released adaptors on non-genetically-modified cells. Towards this end, we purified both ProTag^3^ –comprising an N-terminal 6xHis-tag, sfGFP, a 3xG_4_S linker and a heparin binding domain (HBD, amino acids 123-144 from placenta growth factor-2, PlGF-2)^11^ (**Fig. 3A**)– and a control protein lacking the HBD domain from bacterial culture by high-performance liquid chromatography (**Fig. 3B**). During dialysis of the protein preparations into physiological buffer after purification, the profound contribution of the HBD to protein behavior became apparent: ProTag in a 13 µM stock solution, but not similarly concentrated (9.7 µM) control protein, was found to aggregate in Tris-HCl pH 7.4 (**Fig. 3C**) as well as (to a lower extent) in PBS. Increasing salt concentrations reduced the turbidity measurable after overnight buffer exchange, with dialysis into 1 M NaCl–PBS resulting in no measurable aggregation (**Fig. 3D**).

**Figure 3:**
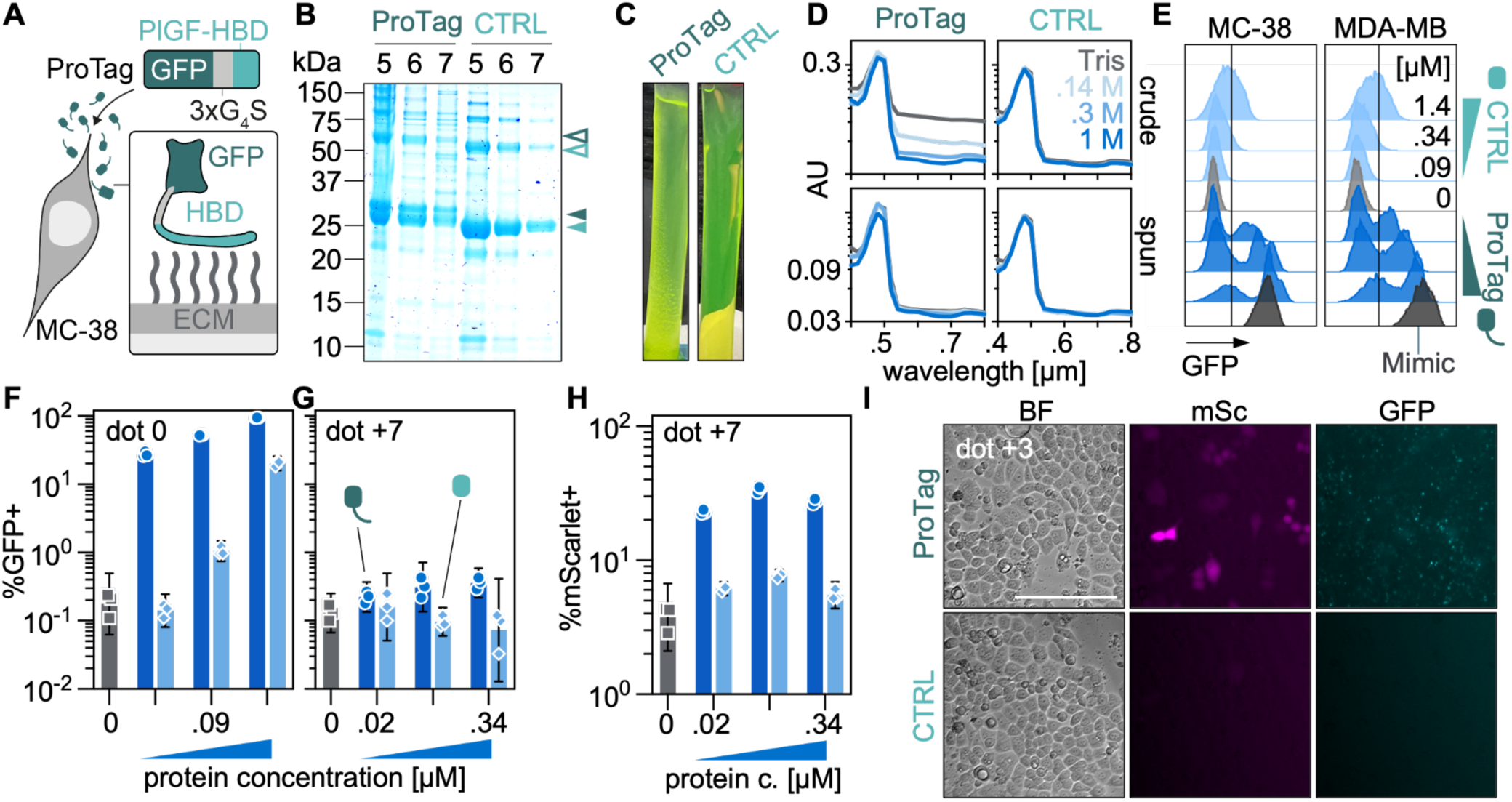
ProTag binds tumor cells and mediates transduction via the PlGF-2 heparin-binding domain. (**A**) The probiotic tag (ProTag) used as an adaptor protein (used here and in a previous study^3^) consists of a superfolder GFP connected via a 3xG_4_S linker to the heparin-binding domain (HBD) of placenta growth factor-2 (PlGF-2), which has been shown to bind extracellular matrix (ECM)^11^. (**B-E**) Assessing HBD-mediated ProTag aggregation and tumor cell-binding. ProTag or a control protein (sfGFP-3xG_4_S, CTRL) lacking the PlGF-2 HDB were purified from *E. coli* via Ni-NTA-HPLC. (**B**) SDS-PAGE of elution fractions (5-7) prior to pooling for ProTag (dark green) and CTRL protein (light green). Solid and empty arrows indicate bands thought to represent monomeric and dimeric ProTag (31.5 kDa) or CTRL protein (28.8 kDa), respectively. (**C**) Photographs of ProTag and CTRL protein solutions in dialysis tubing after over-the-weekend buffer exchange into Tris-HC pH 7.4. Note the turbidity and macroscopic particulates in the ProTag-containing tube. (**D**) Absorption scans to assess turbidity of ProTag- or CTRL-containing samples after overnight buffer exchange into Tris-HCl pH 7.4 or PBS containing 0.14-1 M NaCl, before (crude) or after centrifugation to remove particulates (spun). (**E**) Modally normalized flow cytometry histograms of GFP-signal among singlets after staining of MC-38 and MDA-MB-468 cells with the indicated concentrations of ProTag (dark blue) or CTRL (light blue) protein. Untreated cells (light gray) and GFPMimic cells (dark gray) served as negative and positive controls, respectively. (**F-I**) Transduction of ProTag-decorated cells. (**F**) GFP-positivities on the day of treatment (dot 0) and (**G**) GFP-as well as (**H**) mScarlet (i.e. transgene)-positivities on dot +7 as determined by flow cytometry after treatment of ProTag-(dark blue circles), CTRL-decorated (light blue diamonds) or non-decorated (gray squares) MDA-MB-468 cells with mScarlet-transferring aGFP-LV (1:1 ratio of aGFP-Display and G_mut_). All cell frequencies refer to singlets. Symbols represent data from n=3 technical replicates, bars & error bars represent geometric means ± 95% CIs. (**I**) Fluorescence micrographs of MDA-MB-468 decorated with 0.34 µM ProTag or CTRL, three days after vector treatment (dot +3). Scale bar represents 200 µm.

Crucially, however, the PlGF-2-derived HBD not only mediated self-interaction of ProTag, but was essential for cell decoration: When ProTag or control protein were used as staining reagents for flow cytometry of MC-38 and MDA-MB468 cells, a separation of negative and positive populations was only observed for cells decorated with ProTag, but not with control protein. (**Fig. 3E**). ProTag-treated MDA-MB-468 cells displayed up to 94.7% GFP-positivity and 18.5–fold higher GFP-fluorescence intensities than cells treated with the same concentration of control protein (**Fig. 3F**). This HBD-mediated decoration was transient, with no clearly above-background GFP-signal measured seven days later (**Fig. 3G**). Still, addition of aGFP-LV resulted in 6.0– to 8.9–fold higher mScarlet gene transfer rate on ProTag-decorated cells than on those treated with no protein at all, while transduction rates on cells treated with control protein did not clearly exceed the background rate. The highest mean transfer rate of around 34% was only observed at the second-highest (90 nM), not the highest (340 nM) ProTag concentration tested (**Fig. 3H**). This may be explained by LV particles binding GFP of excess, i.e. non-cell-bound, ProTag and thereby being unavailable for cell transduction, and is in line with the observation of non-cell-bound GFP-positive granulae in ProTag-treated cultures 3 days after decoration & vector addition (**Fig. 3I**).

As the NanoPro system will likely require sustained bacterial tumor presence and continuous release of ProTag for maintenance of the nanovector interface *in vivo*, we sought to maximize protein release without impacting bacterial viability. To this end, we evaluated the effects of deletions targeting three genes in cryptic plasmid-cured *Escherichia coli* Nissle 1917 (EcNc): *lpp*, *lon*, and *ompT*, with the first encoding for the ubiquitous lipoprotein, which tethers the inner and outer membranes of the gram-negative strain used to its peptidoglycan cell wall^12,13^; and the second and third encoding for intra-^14^ and extracellular^15^ proteases, respectively (**Fig. 4A**). When Δ*lpp*, Δ*lon/*Δ*ompT*, and Δ*lpp/*Δ*lon/*Δ*ompT* EcNc were transformed with plasmids encoding either ProTag or control protein, *lon/ompT*-deficient bacteria were found to be at least 200–fold less fit than the parental strain, while no relevant drop in viability was measured for *lpp*-deficient bacteria. Interestingly, the triple-knockout strain appeared more robust than the Δ*lon/*Δ*ompT* strain, displaying only 6.3–fold lower fitness than the parental strain and overlapping 95% confidence intervals with Δ*lpp* readings (**Fig. 4B**). This viability penalty of *lon/ompT*-KO was not a trade-off for higher ProTag release: When GFP release from parental, Δ*lpp*, and Δ*lpp/*Δ*lon/*Δ*ompT* EcNc transformed with lower- or higher-strength constitutive expression plasmids encoding ProTag, control protein or a non-GFP-containing protein after overnight incubation was assessed, no relevant differences in absolute release between Δ*lpp* and triple-mutant strains were identified. Deletion of *lpp* resulted in 36.2– and 16.5– fold higher ProTag release compared to the parental strain for the lower- and higher-strength expression cassettes, respectively, with maximum average concentrations in the supernatant of 15 nM for the lower and 65 nM for the higher strength cassette. (**Fig. 4C**) Depending on the strength of the expression cassette, these supernatant GFP concentrations represented 12.2–16.9% of the total culture signal for Δ*lpp* as opposed to 0.30–0.55% for the parental strain (**Fig. 4D**). Cultures transformed with control protein-encoding plasmid displayed higher rates of absolute and relative release than those transformed with ProTag-encoding plasmid, which might be another consequence of the presence of the PlGF-2 HBD in ProTag. Fixed-volume Western blotting against G_4_S-linkers in pellets and supernatant from Δ*lpp* and parental cultures supported the release patterns observed for GFP fluorescence and indicated the relative homogeneity of produced protein, with the majority of signal appearing at molecular weights expected for ProTag and control protein, and a single lower-weight band possibly representing truncated product (displaying at least 21–fold lower mean gray value than the main band) only observed in pellets from control protein-producing, but not ProTag-producing, bacteria. (**Fig. 4E**) Knockout of *lpp* did not affect the microscopic morphology at endpoint of ProTag-expressing EcNc relative to control-expressing overnight cultures (**Fig. 4F**).

**Figure 4:**
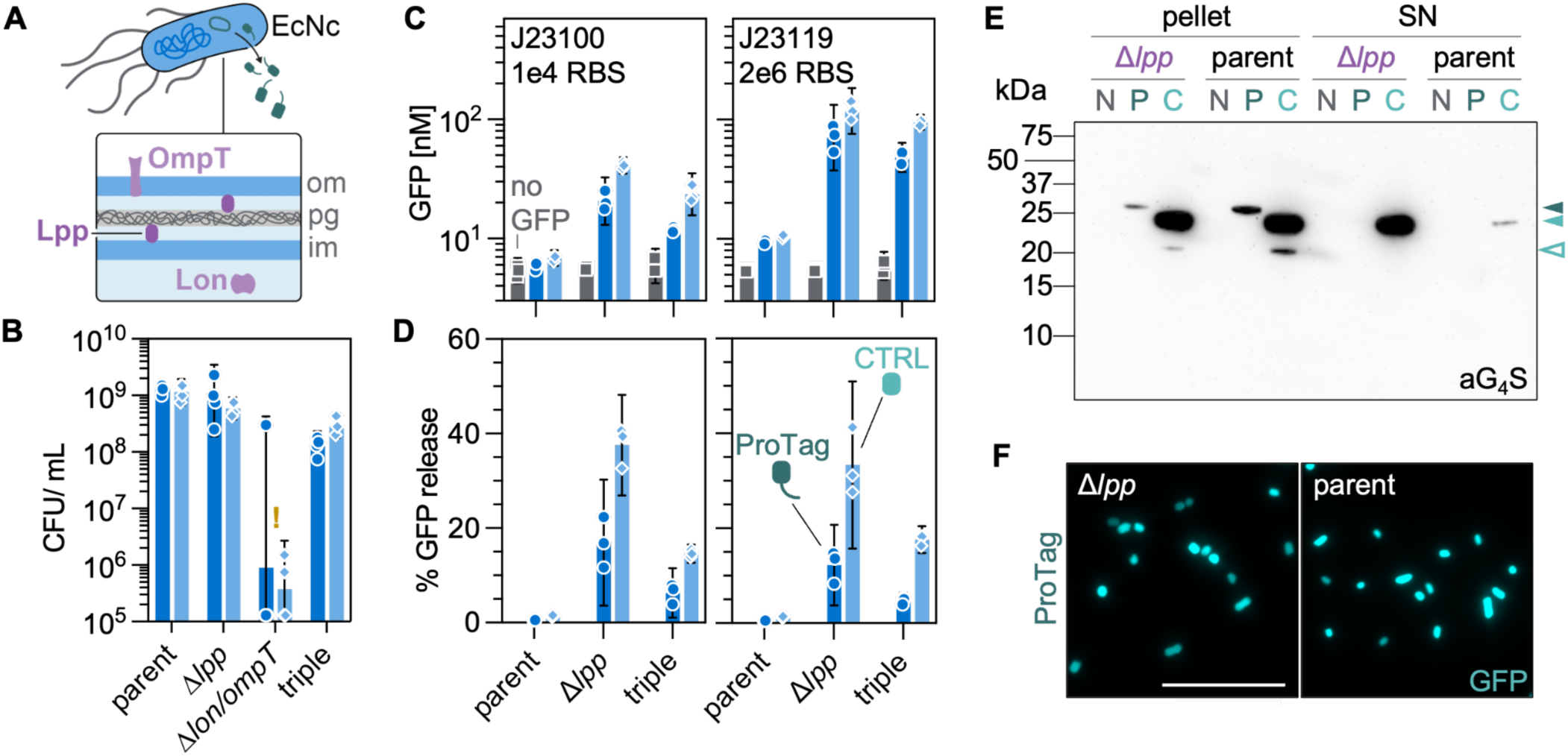
Deletion of *lpp* results in deci-micromolar GFP supernatant release. (**A**) Using an *asd*-deficient *Escherichia coli* Nissle 1917 strain cured for its cryptic plasmids (EcNc) as a chassis, two sets of knockouts were evaluated for their ability to facilitate ProTag release, specifically of: i. *lpp*, which encodes for lipoprotein, a tether of outer membrane (om), inner membrane (im) and peptidoglycan cell wall (pg); and ii. *lon/ompT*, which encode for intra- and extracellular proteases, respectively. (**B**) Assessing the impact of KOs on bacterial viability. Bacterial titer (colony-forming units/ mL) after overnight culture of parental (Δ*asd*), Δ*asd/*Δ*lpp*, Δ*asd/*Δ*lon/*Δ*ompT* or triple KO (Δ*asd/*Δ*lpp/*Δ*lon/*Δ*ompT*) strains transformed with *asd*-stabilized^36^ plasmids encoding ProTag (dark blue circles) or CTRL (light blue diamonds). Note that five clones of ΔasdΔlonΔompT mutant displayed a lawn at one dilution and no colonies at the next lower dilution (denoted with “!” in graph). CFUs were calculated assuming 0.5 observed colonies at the lower dilution. Symbols represent data from n=4 clones, bars & error bars represent geometric means ± 95% CIs. (**C-F**) Characterization of ProTag release. Mutant strains –see (B)– were transformed with *asd*-stabilized plasmids encoding ProTag (dark blue circles), CTRL (light blue diamonds), or a ProTag based on ALFA-Tag as a “no GFP” control (gray squares). Expression cassettes for proteins-of-interest were driven either by a J23100 (iGEM, BBa_J23100) promoter in conjunction with an RBS with a predicted activity^37–42^ for GFPProTag of approx. 1e4 arbitrary units, or by a J23119 (iGEM, BBa_J23119) promoter with a 2e6 AU-RBS (1e5 AU for the ALFAProTag-encoding plasmid). After overnight culture, GFP fluorescence of both crude culture and culture supernatant (obtained by centrifugation) was measured and quantified by comparison with standard curves of purified ProTag and CTRL (see Fig. 3). (**C**) GFP concentration in the culture supernatant, and (**D**) fraction of supernatant-resident GFP signal in comparison to crude cultures. Symbols represent data from n=3 clones, bars & error bars represent geometric means for (C) and arithmetic means for (D), ± 95% CIs. (**E**) Western blot for the presence of G_4_S linkers in supernatants and cell pellets from parental and Δ*asd*Δ*lpp* EcNc overnight cultures transformed with (higher-strength cassette) ProTag- (P), CTRL- (C) or non-GFP (N)-encoding plasmids. Solid arrows indicate bands thought to represent monomeric ProTag or CTRL, empty arrow indicates a fragment of CTRL protein. (**F**) Fluorescence micrographs of parental and Δ*asd*Δ*lpp* EcNc after overnight culture. Scale bars represent 15 µm.

In decoration assays, supernatants from Δ*lpp* EcNc cultures potently decorated mammalian cells and mediated transduction in an HBD-dependent manner (**Fig. 5A**): On the day of decoration, before transduction, both MDA-MB-468 and MC-38 cells displayed above-background GFP signal after treatment with undiluted supernatant or dilutions ranging from 5– to 625–fold. Unlike for the purified protein preparations tested earlier, however, similar GFP-positivities were observed for both ProTag and control protein groups across dilutions. This may be due to the presence of subcellular, GFP-containing bacterial debris which was not cleared by centrifugation and which unspecifically interacted with cells and transiently obscured HBD-mediated ProTag binding. Indeed, HBD-dependent GFP signal was observed for both cell lines four days after treatment. (**Fig. 5BE**) At that time, above-background cytosolic mScarlet lentiviral transfer signal was only present in those cells decorated with ProTag, but not with control protein. As before, transfer rates were highest in those cultures treated with diluted supernatant. Peak mean transfer rates were 52.9% (27.2–fold over background) for 5–fold diluted supernatant on MDA-MB-468 and 49.5% (10.1–fold over background) for 125–fold diluted supernatant on MC-38, respectively, similar to those expected from the same dose of aGFP-LV on genetically GFP-decorated MDA-MB-468 cells, in keeping with the lower LV dose applied to the more permissive MC-38 cells. Notably, above-background transfer signal was observed for both cell lines up to the highest tested supernatant dilution of 625–fold, highlighting the considerable robustness of the established interface. (**Fig. 5 DCF**)

**Figure 5:**
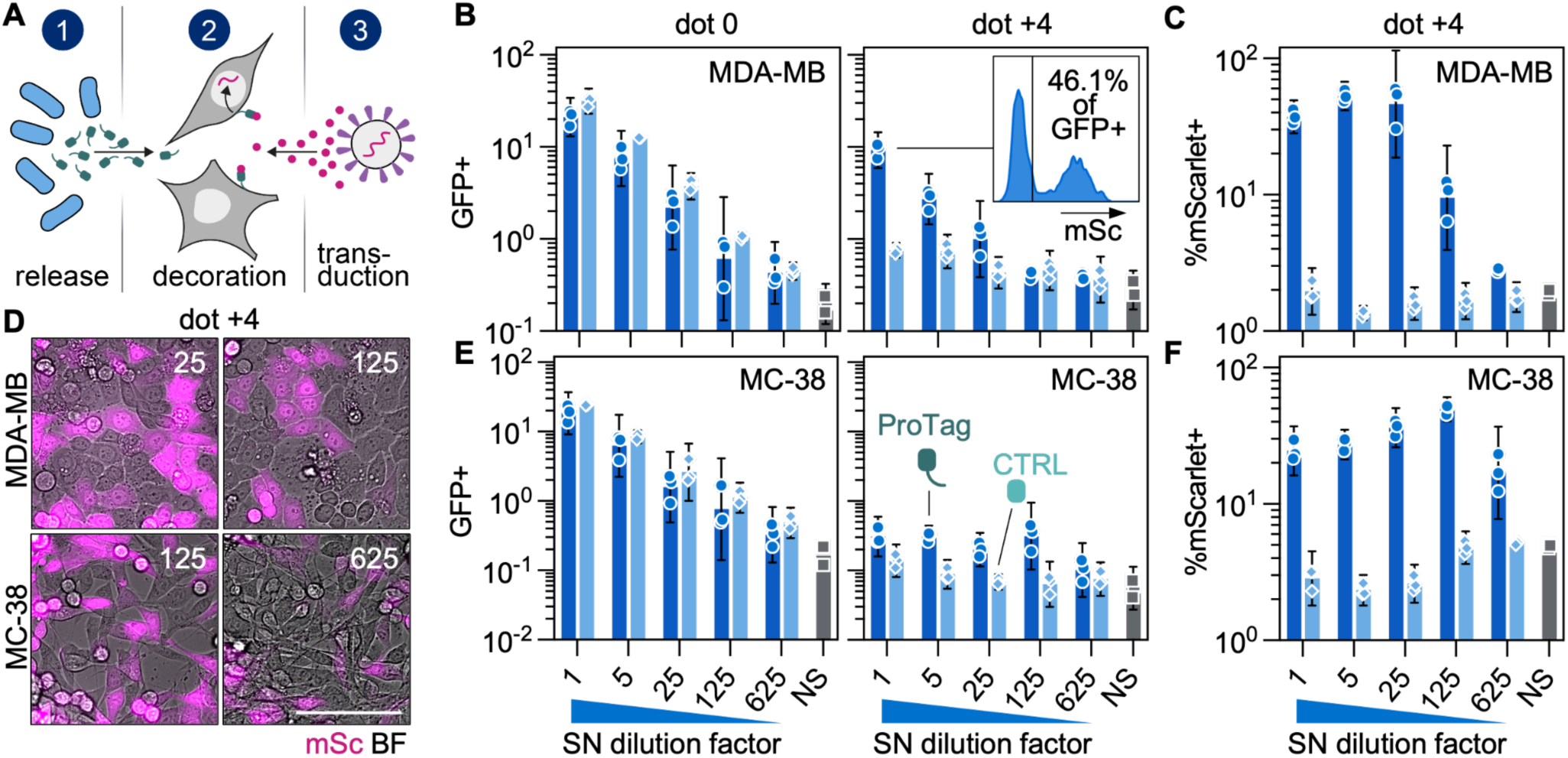
Sub-LOD quantities of bacterially released ProTag enable specific tumor cell conversion. (**A**) Experimental workflow: 1. Supernatants from overnight cultures of Δ*asd*Δ*lpp* EcNc were used to 2. decorate MDA-MB-468 and MC-38 cells, which were subsequently 3. treated with aGFP-LVs. (**BE**) GFP-positivities on the day of treatment (dot 0) and dot +4 as well as (**CF**) mScarlet-positivities on dot +4 as determined by flow cytometry after treatment of MDA-MB-468 or MC-38 cells with the supernatants (or their indicated dilutions) from ProTag-(dark blue circles) or CTRL-producing (light blue diamonds) bacteria or no supernatant (NS) with mScarlet-transferring aGFP-LV. Inlay in (B) shows representative histogram of mScarlet signal distribution on dot +4 among GFP+ cells from a well initially treated with undiluted ProTag-containing supernatant. All cell frequencies refer to singlets. Symbols represent data from tumor cells treated with supernatants from n=3 clones, bars & error bars represent geometric means ± 95% CIs. Error bars were omitted for the left graph of panel E, as one replicate was 0% and therefore unplottable on a logarithmic axis. (**D**) Dot +4 fluorescence micrographs of cells decorated with the indicated dilutions of ProTag-containing supernatant before aGFP-LV treatment, BF: brightfield; mSc: mScarlet. Scale bar represents 100 µm.

Importantly, the sensitivity and specificity of the interface is not restricted to the interaction of GFP and its cognate nanobody, but can be modularly adapted to different antigen:binder-pairs. To demonstrate this, we generated variants of ProTag containing both sfGFP and ALFA-Tag or only the ALFA-Tag (**Fig. 6A**). Contrary to expectations, ALFAProTag was expressed at a lower rate than ALFA-GFPProTag from the same cassette context, with lower ALFA signal in both bacterial pellets and supernatants detected in ELISA (**Fig. 6BC**) –possibly due to impacts of the small size of the ALFAProTag on protein biosynthesis– and above-background ALFA signal on the day of decoration was only detected in cells treated with supernatant from ALFA-GFPProTag-producing bacteria (**Fig. 6D**).

**Figure 6:**
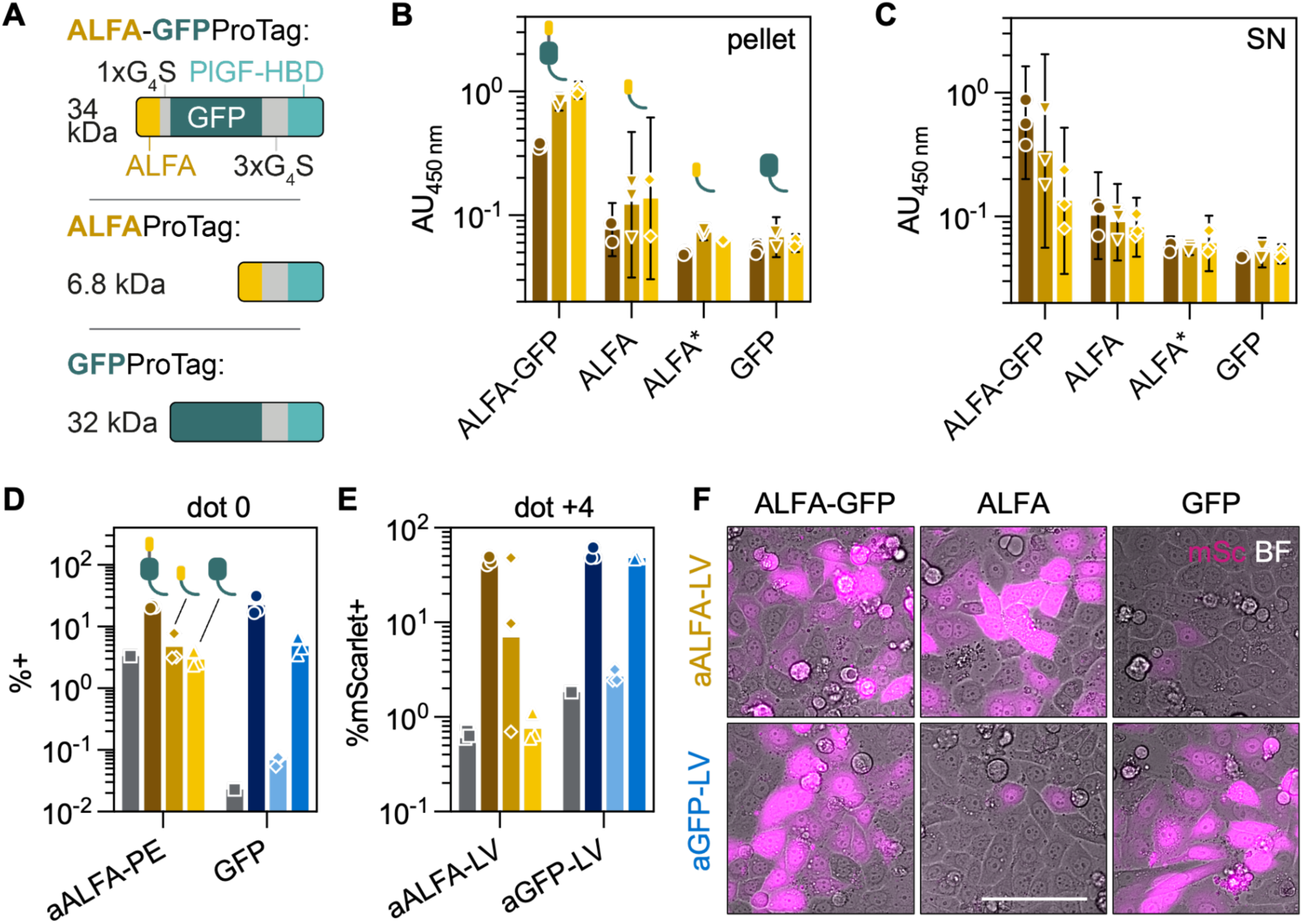
Alternative ProTag architectures for ALFA- and dual-specific orthogonal targeting. (**A**) Architectures and expected molecular weights of the reference GFPProTag as well as ALFA- and ALFA-GFPProTag. (**B-C**) Assessing production & release of ProTag variants. The ALFA-Tag content of (**B**) pellets and (**C**) supernatants from overnight cultures of Δ*asd*Δ*lpp* EcNc transformed with plasmids encoding ALFA-GFPProTag, ALFAProTag or GFPProTag were assessed by ELISA using an aALFA primary antibody and colorimetric readout. All plasmids used the lower-strength expression cassettes specified in Fig. 4C, except the one used to transform the culture marked ALFA*, which carried the higher-strength cassette featuring an ALFA-specific RBS with a predicted activity of approx 1e5 AU. Brown circles: undiluted sample; golden triangles: 10–fold diluted samples; yellow diamonds: 100–fold diluted samples. Symbols represent data from cells treated with supernatants from n=3 clones, bars represent geometric means, error bars 95% CIs. (**D-F**) Comparing ALFA-v. GFP-mediated transduction. MDA-MB-468 cells were decorated as in Fig. 5 with undiluted bacterial supernatant containing ProTag variants (circles: ALFA-GFPProTag; diamonds: ALFAProTag; triangles: GFPProTag) or no supernatant (gray squares), before treatment with either aALFA- or aGFP-LV transferring mScarlet. Frequencies –as determined by flow cytometry– of (**D**) GFP+ as well as ALFA+ events (with the latter determined via an aALFA antibody) on dot 0, as well as (**E**) mScarlet+ events on dot +4. Symbols represent data from cells treated with supernatants from n=3 clones. (**F**) Dot +4 fluorescence micrographs of cells decorated with supernatant containing the indicated ProTag variants before LV treatment, BF: brightfield; mSc: mScarlet. Scale bar represents 100 µm.

Still, four days after cells were treated with ProTag-variant-containing supernatants followed by aALFA- or aGFP-LV, a binary response was observed: above-background mean mScarlet transfer rates of up to 52.5% (69.0–fold over background for aALFA-LV and 28.7–fold over background for aGFP-LV) were only observed in ALFA-decorated cultures inoculated with aALFA-LV and *vice versa*, highlighting the modularity of the targeting approach (**Fig. 6EF**).

## DISCUSSION

The central innovation of the NanoPro system is a probiotically deposited interface of orthogonal adaptors targetable by nanoscale vectors. Our work demonstrates that lentiviral vectors can be effectively targeted to GFP and ALFA-Tag –antigens which do not naturally occur in mammalian hosts– and achieve entry-level selectivities greater than 99%, validating that core tenet. Although adapter-mediated targeting of lentiviral vectors has been demonstrated before^16,17^, these approaches use tandem binders still subject to the antigen problem, modularly targeting cell types via their expression of naturally occurring cell surface-receptors. Via its PlGF-2 HBD, the ProTag utilized here instead binds a wide array of ECM glycoproteins^11^ and thereby cell lines derived from a range of tumors –i.a. of the breast, ovaries, colorectal tract, liver, and pancreas. ProTag decoration is restricted to tumors by the colonization properties of the depositing probiotic^3^, rendering tumor cells antigenically distinct from healthy tissue.

This may enable the tumor-restricted application of modalities such as oncolytic viruses, whose compatibility with the NanoPro interface is supported by ample evidence of the transferability of diverse pseudotypes between the receptor-targeted lentiviral vectors used as model particles here and e.g. vesicular stomatitis and measles viruses^18–20^. Beyond oncolytic viruses, the use of synthetic nanoscale vectors, i.a. mRNA-lipid nanoparticles –which might enable transient, fine-tunable modulation of the solid tumor microenvironment– with the NanoPro interface should be explored. mRNA-LNPs are understood to deliver their payloads via endocytotic processes. Similarly, vesicular stomatitis virus G protein is a type III viral fusion protein known to initiate membrane fusion in a pH-dependent, i.e. endocytosis-dependent, process, suggesting that receptor-targeted LNPs should in principle be compatible with ProTag-mediated targeting. This interface was very robust, with above-background transfer signal still observed after diluting both input vector or bacterial supernatant 625–fold, representing an excellent starting point for the further preclinical development of the NanoPro platform, as *in vivo* testing will likely see only a fraction of the infused doses of both bacterial microscale and (viral or synthetic) nanoscale vectors arrive and interact within the target tumor. Future work towards the system outlined in Fig. 1 should i) optimize bacterial tumor homing and persistence, e.g. reinforcing tumor tropism through biosensors like hypoxia- and lactate-sensing circuits^21^ and (temporarily) attenuate bacterial immunogenicity^22^; ii) address nanovector clearance by the host response, perhaps through the incorporation of anti-phagocytic cues^23^ in the particle architecture and/or the targeted transient inactivation of host immune components^24^; and iii) improve tumor permeability for nanoparticle delivery, harnessing i.a. progress in the electrical^25^ or probiotically-enabled enzymatic^26,27^ disruption of tumor tissue, or by targeting (yet-to-be-identified) transcytosis receptors for bloodstream exit of nanoscale vectors^28^.

While much work remains, our findings demonstrate the principal feasibility of bacterially mediated micro- to nanoscale interaction and provide a framework for collaborative, highly modular therapies.

## MATERIALS & METHODS

### Plasmids

Plasmids encoding VSVG_mut_ (pCMV-Gmut-Amp) envelope and aGFP-Display (pCMV-aGFPdisplay-Amp) protein were based on pMD2.G (Addgene, 12259), linearized by PCR with Q5 polymerase (NEB, M0494S) using primers 5’-CCTGCACAACAGATTCTTCATGTTTGGAC-3’ and 5’-CTCACGTGCTTTGCCAAAGTGATGG-3’. pCMV-aALFAdisplay-Amp was based on pCMV-aGFPdisplay-Amp by digest of the latter with SfiI (NEB, R0123S) and AleI-v2 (NEB, R0685S). Lentiviral transfer plasmids pLV[EF1a-GFP-hCD8a anchor]-Amp, pLV[EF1a-ALFA-hCD8a anchor]-Amp (for generation of ProMimic cells) and pLV[EF1a-mScarlet-T2A-NanoLuc]-Amp (for NanoPro-LVs) were derived from a precursor, 2nd-generation transfer plasmid with a 3’ self-inactivating (SIN) LTR^29^ via digest with SpeI (NEB, R3189S) and NotI (NEB, R3189S). Bacterial constitutive expression plasmids encoding ALFA-GFPProTag (p15A-asd-J23100-His-ALFA-GFPProTag-Kan) as well as lower- and higher-strength cassettes for ALFAProTag (p15A-asd-J23100-His-ALFAProTag-Kan and p15A-asd-J23119-RBShi-ALFAProTag-Kan), GFPProTag (p15A-asd-J23100-His-GFPProTag-Kan and p15A-asd-J23119-RBShi-GFPProTag-Kan), and control protein (p15A-asd-J23100-His-control-Kan and p15A-asd-J23119-RBShi-control-Kan) were based on an *asd*-stabilized p15a backbone, linearized by digest with Nco-HFI/XhoI (NEB, R3193S & R0146S) for ALFA-GFPProTag, AflII/XbaI (NEB, R0520S & R0145S) for lower-strength ALFAProTag, and BamHI-HF/XbaI (NEB, R3136S & R0145S) for all others. The insert for p15A-asd-J23100-His-GFPProTag-Kan was obtained by BamHI-HF/XbaI digest of the ProTag used by Vincent et al.^3^; all other inserts were ordered as dsDNA fragments (Twist Bioscience). Throughout, NEBuilder HiFi DNA Assembly Mix (NEB, M5520AA) was used for Gibson assembly. Lentiviral envelope and transfer plasmids were amplified in Stbl3 *E. coli* (Thermo, C737303), bacterial plasmids in NEB 5-alpha *E. coli* (NEB, C2987). Bacteria were cultured on Lennox agar plates (Invitrogen, 22700-025) or shaken at 245 rpm in LB Broth Miller (Fisher, BP9723-500), both at 37°C and containing either 100 µg/mL ampicillin sodium (Sigma-Aldrich, A9518-25G) or 50 µg/mL kanamycin sulfate (Fisher, BP906-5). Cryostocks were deposited by mixing one volume of overnight culture and one volume of 50% v/v glycerol (Fisher, BP229-1) in water and stored at -80°C. Plasmid DNA was isolated using QIAprep Spin (Qiagen, 27104) and NucleoBond Xtra Midi Plus (Macherey-Nagel, 740412.50) kits for mini and midi scales, respectively. Identity of each prepared plasmid stock was confirmed by whole plasmid-sequencing (Plasmidsaurus) before use.

### Probiotic Strains

*Escherichia coli* Nissle 1917, cured of their cryptic pMUT1 and pMUT2 plasmids^30,31^ (EcNc), were the basis of all engineered probiotic strains described here and harbour an integrated *luxCDABE* cassette within the genome, which also contains an erythromycin resistance gene^32^. EcNc Δ*asd* were generated by replacing the *asd* gene with a selection marker using the pKD3 system, followed by FLP-mediated recombination^33^. EcNc Δ*asd*Δ*lpp* were then derived from them by pCas/pTargetF-mediated^34^ deletion of *lpp*. EcNc Δ*asd*Δ*lon*Δ*ompT* were generated using the pKD3 system, leaving a chloramphenicol-resistance gene in place of *ompT*. The Δ*asd*Δ*lpp*Δ*lon*Δ*ompT* (“triple-KO”) strain was then engineered from it by deleting *lpp* as described above. All strains described in this study were cured of the helper plasmids introduced for engineering before use. Strain identities were confirmed by sequencing for all except EcNc Δ*asd*Δ*lon*Δ*ompT*, for which colony PCR was employed. Probiotics were rendered electrocompetent by four washes with ice-cold Milli-Q water (Millipore; 3,000 x *g*, 10 min, 4°C) and then stored in water containing 15% glycerol in 100 µL aliquots of approximately 4 ODmL each. Bacteria were transformed with plasmids-of-interest by electroporation using a micropulser (program “EC1”; Bio-Rad) and 0.1 cm gap cuvettes (Bio-Rad, 165-2089).

### Mammalian cells

Lenti-X 293T cells (TaKaRa, 632180) were maintained in DMEM/F12, GlutaMAX (Gibco, 10565-018) supplemented with 9% v/v fetal bovine serum (FBS, Gibco, A52567-01); MDA-MB-468 cells (ATCC, HTB-132) and their ProMimic derivatives (see below) in DMEM/F12, GlutaMAX with 9.1% v/v FBS and 0.9% v/v (i.e. 1X) penicillin/streptomycin (PS, Gibco, 15070-063); MC-38 cells (Kerafast, ENH204-FP) in MC-38 media –DMEM/F12, GlutaMAX supplemented with 9% v/v FBS, 0.9% v/v PS, 0.5 mM sodium pyruvate (Gibco, 11360-070), 10 mM HEPES (Gibco, 15630-080), and 1X non-essential amino acids (NEAA, Gibco, 11140-050), all following suppliers’ specifications regarding density. For subculture, cells were incubated in TrypLE Express detachment reagent (Gibco, 12605-010) for 5 min (MC-38, Lenti-X 293T) or 15-20 min (MDA-MB-468) at 37°C before quenching with 1 to 1.4 volumes of growth media. MDA-MB-468 cells were additionally washed with one volume of TrypLE prior to detachment. For cryopreservation, cells were resuspended at densities between 1e5 and 1e7 cells/mL in the respective culture medium + 5% DMSO (Fisher, BP231-100) before transfer to -80°C in CoolCell (Corning, 432000), followed by long-term storage in the gas phase above liquid nitrogen as soon as 24 h later. To generate ALFA- and GFPMimic cells, MDA-MB-468 cells were transduced with VSVG-LVs (see below) transferring cassettes encoding ALFA or GFP fused to a human CD8a-derived membrane anchor. Cells were treated with series of 5–fold dilutions of each vector stock. Seven days after vector addition, cells were detached (see above) and GFP/ALFA-positivity was determined using an Attune NxT acoustic focusing flow cytometer (Invitrogen). Prior to measurement, ALFA-Mimic candidates were stained (30 min, RT, in the dark) for the presence of ALFA using 25 nM FluoTag-X2 antiALFA Atto488 (NanoTag Biotechnologies, N1502-At488-L). Cultures displaying <30–50% positivity were chosen to obtain material with <5 vector copies/well^35^; transgene-positive cells were selected on an LE-MA900FP multi-application sorter (Sony) 3 days later to deposit polyclonal ALFAMimic and GFPMimic cultures.

### Lentiviral vectors

Lenti-X 293 T cells (see above) –seeded at 1.2e5 cells/cm^2^ the day before, usually in 150 mm dishes– were transfected with envelope plasmid, transfer plasmid, and the packaging plasmid psPAX2 (Addgene, 12260) in a 35:100:65 ratio (200 ng/cm^2^) using NovaCHOice transfection and booster reagent (Sigma-Aldrich, 72622-M) at ratios of 1 µL/ng DNA and 0.5 µL/ng DNA, respectively, with complex formation taking place in plain DMEM/F12, GlutaMAX. The sole envelope plasmid for production of VSVG-LVs was pMD2.G (see above); for NanoPro-LVs, pCMV-Gmut-Amp and pCMV-aALFAdisplay-Amp or pCMV-aGFPdisplay-Amp were used at the indicated weight ratios for a total of 35 ng/cm^2^ of envelope plasmid DNA. Plasmid-containing supernatant was replaced with growth medium 6-16 h post addition. Supernatants were then harvested 36-48 h after medium exchange and clarified by clarification through 0.45 µm PVDF syringe-tip filters (Merck Millipore, SLHVR33S). Three volumes of clarified supernatant were mixed with one volume of Lenti-X concentration reagent (TaKaRa, 631231) and incubated at 4°C overnight before centrifugation (1,500 x *g*, 4°C, 45 min) and 100–fold concentration of lentiviral particles by resuspension of the pellet into DPBS (Gibco, 14190-144). Concentrated stocks were aliquoted and stored at -80°C.

### Transducing ProMimic cells

One day before vector treatment, monocultures of MDA-MB-468, ALFA-, or GFPMimic cells were seeded at 5e3 – 6.4e3 cells/96-well; 1:1 cocultures of ALFA- and GFPMimic cells were seeded at a total density of 1.9e5 (Fig. 2G) – 4e5 cells/6-well (Fig. 2H). After attachment, cells were treated with NanoPro-LVs at the indicated doses diluted in growth medium, for total volumes of 200 µL/96-well and 3 mL/6-well. Three (Fig. 2GH) to four (Fig. 2IJ) days after vector treatment, well contents were analyzed by epifluorescence microscopy (Ti2, Nikon) and flow cytometry (Attune NxT, Invitrogen).

### Purification of ProTag & control protein

NEB 5-alpha *E. coli* were transformed with Axe/Txe-stabilized plasmids encoding either N-terminally His-tagged ProTag^3^ (see above) or a control protein-encoding plasmid generated from it by truncating PCR (using primers 5’-TAATCTAGAACTAGTCTTGGACTCCTGTTGATAGATCCAG-3’ and 5’-GGAACCCCCGCCACCC-3’) followed by ligation with KLD enzyme mix & buffer (NEB, M0554S & B0554A). Single clones were used to inoculate 2 mL overnight cultures, which were then used to inoculate 1 L cultures. These were incubated at 37°C, 225 rpm for 9 h, followed by 18 h at 25°C. Bacteria were pelleted and frozen at -80°C when cultures had reached OD600 of 3.2 and 4.3, respectively. Later, pellets were thawed and resuspended in lysis buffer (50 mM NaH_2_PO_4_–300 mM NaCl, pH 8.0) on ice. DNAse and RNAse were added to suspensions to final concentrations of 10 µg/mL each before cells were lysed by sonication (40% amplitude, 30 min, 2 s pulse – 4 s rest). Lysates were cleared by centrifugation (20,000 x *g*, 30 min, 4°C) before His-tagged proteins were purified by liquid chromatography on a ÄKTA FPLC system (Cytiva) using a 25 mL HisPur™ Ni-NTA resin column (Thermo Scientific) using 50 mM NaH_2_PO_4_–1 M NaCl–20 mM imidazole, pH 8.0 wash buffer and 50 mM NaH_2_PO_4_–1 M NaCl–250 mM imidazole, pH 8.0 elution buffer. Eluate fractions were analyzed by SDS-PAGE on a 10% Bis-Tris gel (Invitrogen, NW00105); notably, sample buffer did not contain DTT. Optimal storage conditions were assessed in a dialysis trial: 500 µL/ tube of ProTag or control protein were loaded into 0.5 mL dialysis devices with 3.5 kDa molecular weight cut-off (Thermo, 88400) together with 14 mL/tube of Tris-HCl or the indicated DPBS variants. After overnight incubation at 4°C, absorption scans (400 – 800 nm) of buffer-exchanged material before and supernatant after centrifugation (20,000 x *g*, 5 min, 4°C) were performed on an infinite 200Pro plate reader (Tecan). Based on trial results, material was loaded into 3.5 kDa molecular weight cut-off dialysis tubing (Spectrum Labs, 132724) and incubated in 1 M NaCl – DPBS overnight. Dialyzed material was then frozen at -80°C after addition of glycerol to a final concentration of 20% v/v. Concentration of purified and dialyzed protein was determined spectrophotometrically by measuring absorption at 488 nm, assuming ε = 8.33e4.

### Supernatant release assays

To assess the impact of deletions on supernatant release, the relevant strains were transformed with ProTag-encoding plasmids (as described above) and initial overnight cultures in 96-well deepwell plates were used to inoculate a second culture plate at an OD of 0.01 for overnight incubation. The next day, optical density at 600 nm (OD_600_) as well as GFP fluorescence (Ex: 488 nm; Em: 520 nm) of both crude cultures and supernatants (after centrifugation at 3,000 x *g*, 10 min, RT) were determined using an infinite 200Pro plate reader (Tecan). A standard curve of purified ProTag and control protein in LB broth (see above) was used to quantify GFP release. To determine impact of mutations on bacterial viability, 1:10 dilution series (with dilution factors ranging from 1e1 – 1e8) of cultures transformed with (lower-strength) ProTag and control protein-encoding cassettes were prepared and plated onto square kanamycin-containing LB agar plates at 4 µL/well. The next day, colonies per drop & dilution were counted to determine colony-forming units/mL. Size distribution of released protein was assessed by Western blot analysis: Approx. 1.6 – 2.4 ODmL/sample of crude overnight cultures (inoculated directly from glycerol stocks) as well as equivalent volumes of culture supernatant were collected. Bacteria were pelleted and lysed using BugBuster protein extraction reagent (Millipore, 70584-3), and both pellet and SN samples –diluted 1:2 in sample buffer containing DTT at a final concentration of 0.5 M– were run on a 12% NuPAGE Bis-Tris gel (Invitrogen, NP0342BOX) at 100 V for 40 min followed by 45 min at 150 V. The gel was washed with water prior to transfer onto a nitrocellulose membrane (Invitrogen, IB23002) using an iBlot2 system (Invitrogen). After transfer, the membrane was blocked for 40 min with 2% w/v milk–PBS Tween before addition of a rabbit–anti–G4S primary antibody (Genscript, A02310) to a final concentration of 1:1e4 and incubation for 30 min at RT. Following 5 washes (1, 5, 1, 1, 5 min) in PBS-Tween, the membrane was incubated with a secondary, donkey–anti–rabbit, HRP-conjugated antibody (Fisher, 711-035-152; 1:2.5e4 in blocking buffer, see above) for 30 min at RT. After another set of five washes, signals were detected using Immobilon Forte HRP substrate (Millipore, WBLUF0100) on a ChemiDoc imager (Bio-Rad).

To assess production & release of ALFA-containing protein, ELISA was performed on supernatants and pellets from 2 ODmL per overnight culture (inoculated directly from agar plates) in PBS. Pellets were lysed via BugBuster supplemented with 1:1e3 rLysozyme (Millipore, 71110-1200KU) and 1:1e3 Benzonase (Millipore, 71205-3) for 10 min at RT. Samples were used to coat untreated 96-wells (Corning, 3631) for 30 min at RT. After blocking with 1% BSA–PBS-Tween (10 min, RT) and washing with PBS-Tween, primary rabbit–anti–ALFA-Tag antibody (NanoTag Biotechnologies, N1581) was added to a final concentration of 50 ng/µL in 0.1% BSA–PBS-Tween & incubated, 30 min at RT. Wells were washed four times with PBS-Tween before secondary incubation with donkey–anti–rabbit, HRP-conjugated antibody (Fisher, 711-035-152) diluted 1:2e4 in 0.1% BSA–PBS-Tween (30 min, RT). Wells were then washed four times with PBS-Tween before induction of a colorimetric reaction with Turbo TMB-ELISA solution (Thermo Scientific, 34022), stopped with 2 M sulfuric acid before measurement on an infinite 200Pro microplate reader (Tecan).

### Decoration assays with purified protein

Cells were detached from 150 mm dishes by scraping with 1.8 cm blades (Fisher, 08-100-241) and singularized by passage through 70 µm strainers (Corning, 431751) and pipetting. Cells were then distributed to V-bottom 96-wells at approx. 1e5 cells/well and treated with ProTag or control protein diluted in stain buffer (BD Pharmingen, 554657; Fig. 3E) or LB broth (Fig. 3FGH): material was centrifuged (400 x *g*, RT, 5 min), supernatant was removed, pellets were resuspended in 45 – 75 µL/well of protein dilution, and samples were incubated at 37°C for 30 min (Fig. 3FGH) or at RT for 50 min (Fig. 3E), in the dark. After removal of protein dilutions and resuspension, the material was distributed for flow cytometric analysis and transduction (in flat-bottom 96-well plates), the latter using 0.9 µL/well of aGFP-LV diluted in growth medium for a total culture volume of 200 µL/well. Three days after treatment, well contents were documented by epifluorescence microscopy and subcultured 1:2 into fresh 96-well plates. Flow cytometric analysis was performed seven days post treatment (dpt).

### Decoration assays with bacterial supernatant

MDA-MB-468 and MC-38 cells were decorated with bacterial supernatants much like described for the purified protein above. After detachment and singularization, 2.5e4/96-well and 1.3e4/96-well, respectively, MDA-MB-468 and MC-38 cells were incubated in 75 µL/well of bacterial supernatant –diluted in growth medium, where applicable– for 30 min at 37°C in the dark. Following decoration, 5e3 MDA-MB-468 or 1.3e3 MC-38 cells each were seeded into flat-bottom 96-wells and 0.5 or 0.1 µL/well of aGFP-LV diluted in growth medium were added, respectively, to a final volume of 200 µL/well. aALFA-LV was used on MDA-MB-468 cells at 0.6 µL/well. Flow cytometric and epifluorescence microscopic analyses were performed 4 dpt. On dot 0, the non-seeded decorated cell material was stained for the presence of ALFA-Tag using a polyclonal rabbit–anti– ALFA-Tag primary antibody (NanoTag Biotechnologies, N1581) at 0.33 µg/mL and a donkey–anti–rabbit IgG, PE-conjugated secondary antibody at 0.08 µg/mL, before both ALFA- and GFP-decoration were assessed cytometrically.

## ACKNOWLEDGEMENTS

The authors thank Dorothee M. Günther (New York University, New York, USA) as well as Christian J. Buchholz and the members of his lab (Paul-Ehrlich-Institut, Langen, Germany) for fruitful discussions about experiment and vector design as well as data analysis. This work was supported by the National Institutes of Health, under grants R01EB030352 and R01CA313939, both held by Tal Danino.

## AUTHOR CONTRIBUTIONS

**Conceptualization:** AM, ZSS, ERB

**Methodology:** AM, ERB

**Investigation:** AM, AH, MA, AML, KRG, AL, ERB

**Formal Analysis:** AM

**Resources:** RLV, ERB, JI, AML, AL

**Visualization:** AM

**Writing – original draft:** AM, MA

**Funding Acquisition:** TD

**Supervision:** AM, TD

